# 3D-Scaffold: Deep Learning Framework to Generate 3D Coordinates of Drug-like Molecules with Desired Scaffolds

**DOI:** 10.1101/2021.06.02.446845

**Authors:** Rajendra P. Joshi, Niklas W. A. Gebauer, Mridula Bontha, Mercedeh Khazaieli, Rhema M. James, Ben Brown, Neeraj Kumar

## Abstract

The prerequisite of therapeutic drug design is to identify novel molecules with desired biophysical and biochemical properties. Deep generative models have demonstrated their ability to find such molecules by exploring a huge chemical space efficiently. An effective way to obtain molecules with desired target properties is the preservation of critical scaffolds in the generation process. To this end, we propose a domain aware generative framework called 3D-Scaffold that takes 3D coordinates of the desired scaffold as an input and generates 3D coordinates of novel therapeutic candidates as an output while always preserving the desired scaffolds in generated structures. We show that our framework generates predominantly valid, unique, novel, and experimentally synthesizable molecules that have drug-like properties similar to the molecules in the training set. Using domain specific datasets, we generate covalent and non-covalent antiviral inhibitors. To measure the success of our framework in generating therapeutic candidates, generated structures were subjected to high throughput virtual screening via docking simulations, which shows favorable interaction against SARS-CoV-2 main protease and non-structural protein endoribonuclease (NSP15) targets. Most importantly, our model performs well with relatively small volumes of training data and generalizes to new scaffolds, making it applicable to other domains.

## Introduction

The COVID-19 pandemic, caused by SARS-CoV-2, posed a serious challenge to the public health worldwide.^1^ With the aim to address such challenges in developing lead candidates against different diseases, it is necessary to have a disease aware generative model that quickly generates effective therapeutics from the unknown and massive chemical space and could be tested with cell based assay screening.

The discovery and development of a new therapeutic is a long, expensive, and risky process that sometime takes many years before clinical approval. One of the challenges in drug design is to find small molecules with desired functionalities. ^2^ This is a daunting task with conventional methods, which has slowed down the discovery of high impact molecules for diverse applications. ^3^ The huge chemical space (10^60^) of molecules still remains unexplored.^4–6^ Recently, with the rise of deep learning models, several approaches to efficiently explore the astronomically large chemical space have been proposed. The majority of existing approaches focus mainly on *de-novo* drug design using variational auto-encoders, generative adversarial networks, or reinforcement learning generating molecules mainly in the form of SMILES strings.^7–18^

An alternate and robust way to find drug-like molecules is by generating molecules with desired functional groups, core structures, or scaffolds. ^19,20^ Such fragments play an important role in determining the functionalities of generated molecules, thus making tuning of properties more flexible. Moreover, molecules with certain scaffolds are likely to have desired interactions with a given protein target as a drug. Scaffold-based approaches allow to incorporate such prior knowledge in the generation process in order to increase the chances of obtaining molecules with desired properties compared to simply generating molecules from scratch. Several approaches have been proposed recently to generate therapeutic candidates building on the core structures. ^20–23^ Some of these methods are constrained to certain definitions of scaffolds (e.g. Murcko^24^ scaffolds) or do not guarantee that the desired scaffold is always preserved during molecule generation while others do not generalize well for new scaffolds. ^21,23^ To the best of our knowledge, none of the existing approaches focuses on generating 3D coordinates of drug-like molecules that can be directly tested against the protein target via computational and experimental screening. However, 3D coordinates of generated molecules are required for physics-based simulations as well as for robust graph-based predictive models for modeling the properties. Moreover, these molecules can be directly used for high throughput virtual screening through structure based docking against the proteins to determine their affinity and efficacy as drugs against a particular disease. Consequently, we believe that 3D molecule generation will accelerate the hit identification and lead optimization for drug discovery and development.

In this work, we propose a deep learning framework called 3D-Scaffold that can generate 3D coordinates of therapeutic candidates given a desired scaffold. It is guaranteed that 100% of the generated molecules contain the desired scaffold and the model generalizes well to previously unknown scaffolds not included in the training data. Our current framework is different from existing scaffold-based approaches for multiple reasons: (I) In contrast to existing approaches, which generate SMILES strings or molecular graphs, our model generates 3D coordinates of the molecules with a given core structure; (II) It works equally well for any type of scaffold definition including BM, cyclic skeletons, or side chains provided SMILES strings exist for the desired scaffolds; (III) Our model is transferable to generate molecules with novel scaffolds where the model is not trained on; (IV) Without constraining the model directly on desired properties, our model can generate molecules with properties similar to the training set.

A few issues arise when constructing physics informed machine learning approaches based on 3D nuclear coordinates in contrast to more abstract molecule representations such as strings or molecular graphs. ^25^ The coordinate representation is not invariant to rotation, translation, and indexing of atoms while most properties of interest (e.g. the potential energy or the logP score) are invariant to these transformations or change equivariantly (e.g. atomic forces rotate and translate with the coordinates). The G-SchNet^26^ neural network architecture used in our 3D-Scaffold framework systematically obeys these constraints by design. This allow our model to extract features from the coordinates that capture local symmetries and are invariant to rotation, translation, and indexing of the input coordinates. Furthermore, the distributions it predicts for atom positions equivariantly rotate and translate with respect to the coordinates. Most importantly, we show that our framework designs reasonable molecules even with small training datasets due to the robust architecture of the underlying model. By training it on limited, already known and drug-like molecules, we aim to generate more and previously unseen novel candidates with desired scaffolds that can be synthesized, which ultimately will contribute towards accelerating the discovery of therapeutic drugs.

In this contribution, we applied our 3D-Scaffold for de-novo discovery of molecules specifically tailored to bind with given SARS-Cov-2 diseases targets. Our methodology is exemplified by the task of designing antiviral candidates to target SARS-CoV-2 related proteins. Using carefully curated covalent and non-covalent antiviral datasets, we were able to constrain the generation space for domain aware deep generative framework to generate novel covalent and non-covalent inhibitor candidates. Properties of generated molecules are com-pared with the molecules in the training set. Generated 3D-coordinates of molecules were further examined for their efficacy as antiviral inhibitors against SARS-COV-2 main protease (Mpro) and a SARS-CoV-2 non-structural protein endoribonuclease (NSP15).

## Methods

### 3D-Scaffold framework

The 3D-Scaffold framework is based on an autoregressive, generative deep neural network named G-SchNet,^26,27^ which builds molecular structures from scratch by iteratively placing one atom after another in 3D space, respecting global and local symmetries by design. The neural network uses SchNet for feature extraction,^28–31^ a state-of-the-art predictive model that can predict several quantum mechanical properties of small molecules with benchmark chemical accuracy. In 3D-Scaffold, instead of starting from scratch, molecules are build around a desired scaffold.

From a computational perspective, the neural network used in our framework for *de-novo* therapeutic candidate design is broken down into two major blocks: feature learning and atom placement as shown in Figure 1. In the feature learning block, embedding and interaction layers are used to extract and update rotationally and translationally invariant atom-wise features that capture the chemical environment of an unfinished molecule. Here, the neural network utilizes continuous-filter convolution layers as a means to learn robust representations of molecules starting only from positions of atoms and corresponding nuclear charges. In the atom placement block, the extracted features are used to predict distributions for the type of next atom and its 3D coordinates, where the latter distribution is constructed from predictions of pairwise distances between the next atom and all preceding atoms. In order to do the actual placement of the next atom in 3D space, a distribution on a small grid with candidate positions focused on one of the preceding atoms is constructed from the predicted pairwise distances. The whole procedure is repeated successively in order to build a complete molecule with desired scaffold: after the type and position of the next atom has been sampled from the predicted distributions, new atom-wise features incorporating the added atom are extracted in the feature learning block and then used to place the following atom with the atom placement block.

**Figure 1:**
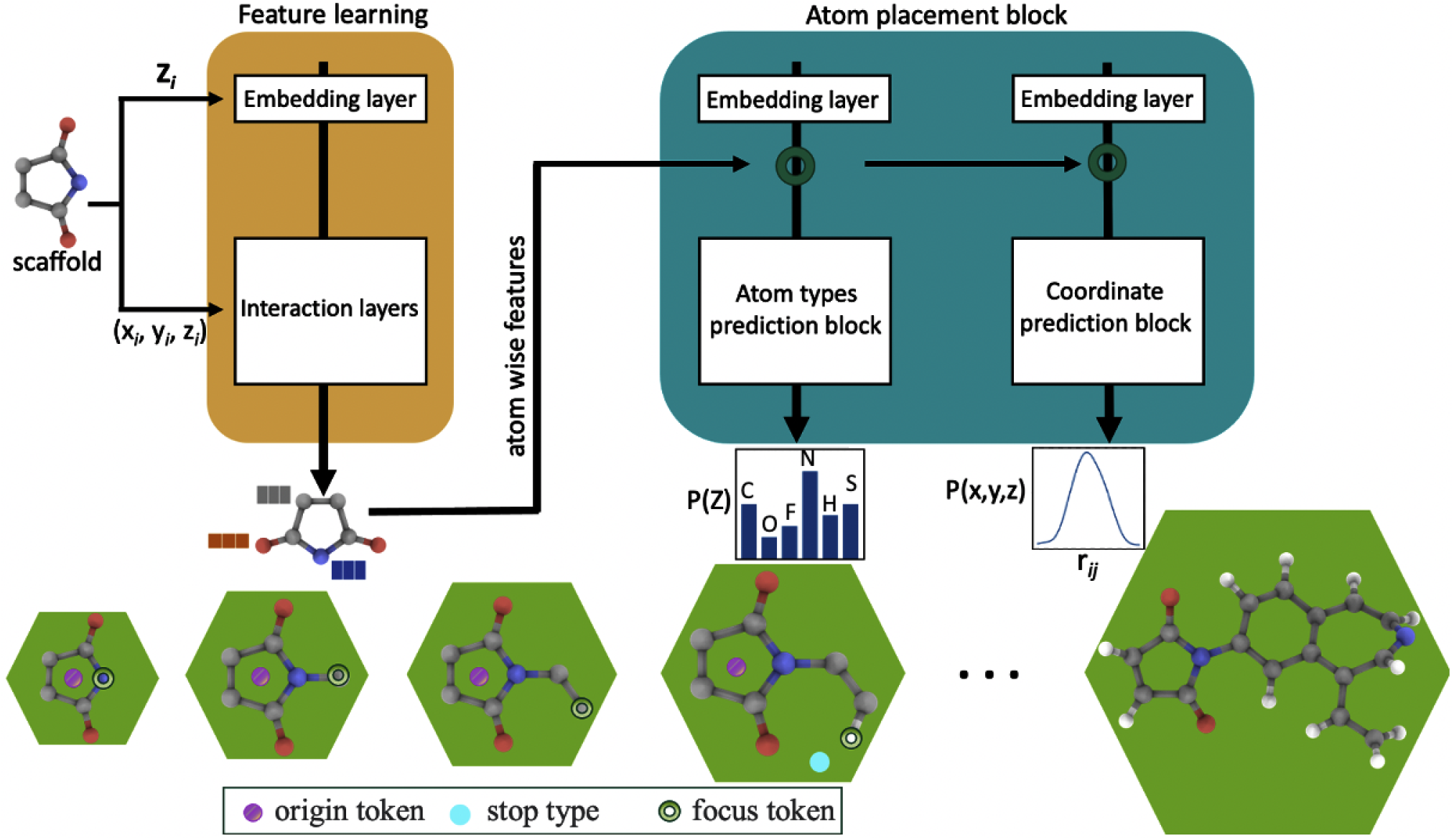
3D-Scaffold framework used as generative model to produce drug-like molecules with desired functionality. The bottom panel shows the scaffold-based molecular generation scheme, where origin token, focus token, and stop type aid the generation of the molecules from scaffolds.

The generation process is aided by two auxiliary tokens with unique, artificial types, namely the origin and focus tokens. At each generation step, one of the already placed atoms is uniformly randomly chosen as focus token. The origin token, in contrast, stays fixed throughout the whole generation procedure. In previous work with G-SchNet by Gebauer et al. ^26^, the origin token marks the center of mass of molecules. In our 3D-Scaffold framework, however, we instead use it to mark the center of mass of the scaffold that is the starting point of the generation procedure. At each step, the unplaced neighbour of the focus token that is closest to the origin token is supposed to be sampled. This means that while the structure grows around the center of mass of the resulting molecule in the previous G-SchNet model, in our current 3D-Scaffold framework it grows from the center of mass of the desired scaffold given to the model as a starting point. If the currently focused atom has no neighbors left to place, the model should predict the stop type instead of a proper atom type and in this way mark the focused atom as finished. Atoms marked as finished cannot be chosen as focus anymore and after all atoms have been marked as finished, the generation process terminates. The resulting schemes for training of the model and generation of molecules are summarized as pseudo code in Table 1.

**Table 1:**
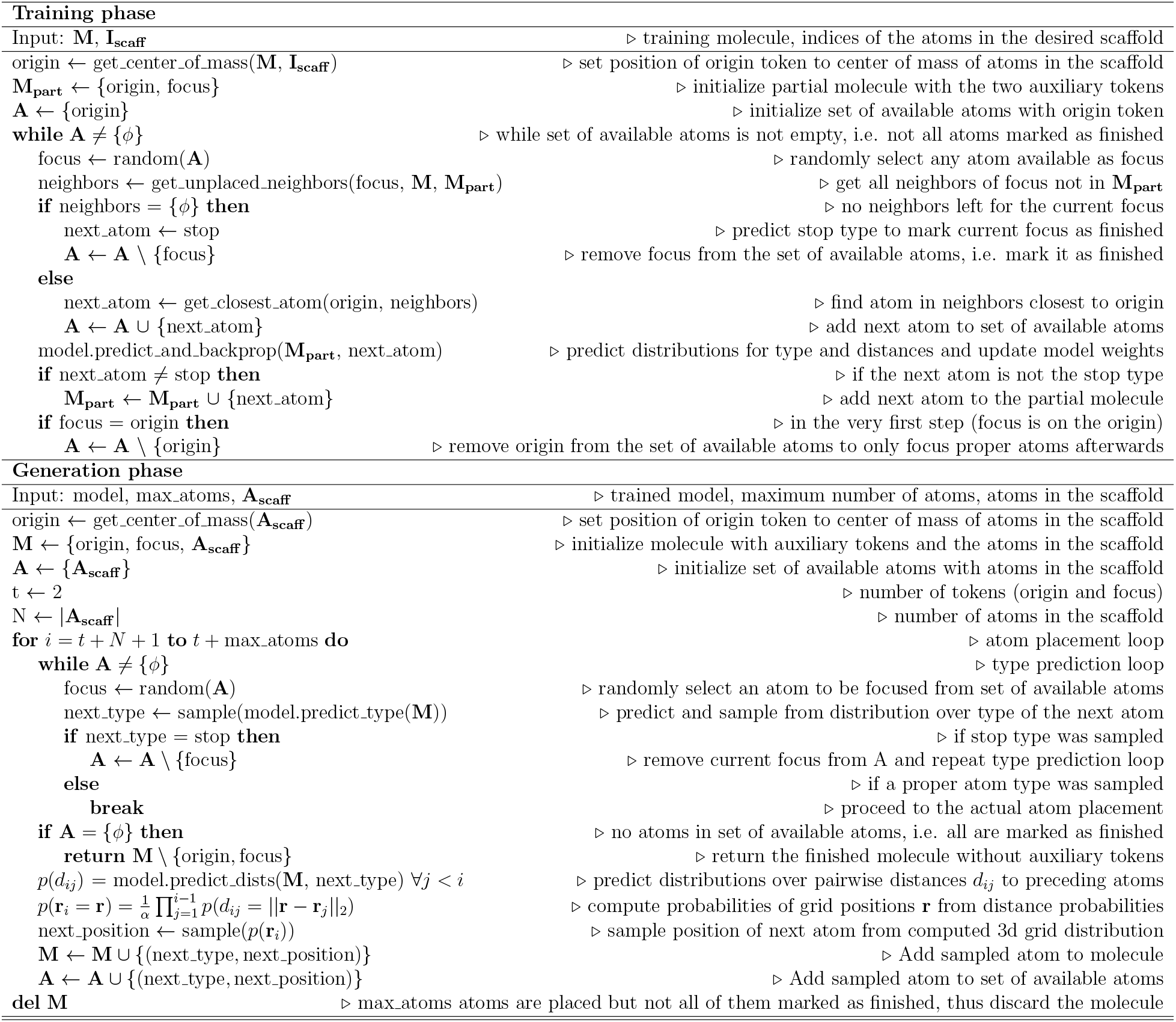
Pseudo code for training and generation phases in 3D-Scaffold framework.

The model is trained end-to-end with backpropagation using the ground truth types and pairwise distances of atoms in training data molecules split into sequential atom placement steps as described in the pseudo code. At each training step, the model predicts the type of the next atom and its distances to all preceding atoms. The distributions predicted by the model are discrete: the type distribution contains a probability value for each atom type occurring in the training dataset and the stop type and the distance distributions cover distances between 0 *Å* and 15 *Å* in 300 equally spaced bins. At any step, let *Z*_next_ be the ground truth type of the next atom and 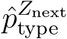 the probability that the model assigns to that type at the current step. Then, we use negative log-likelihood as the loss for the type prediction:

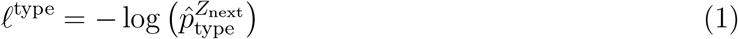

For the loss on distance predictions, we use the cross-entropy between true and predicted distances:

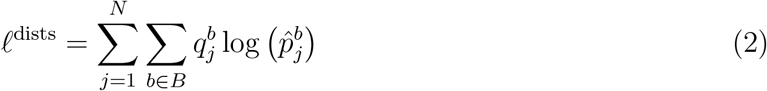

with Gaussian expanded ground truth distances

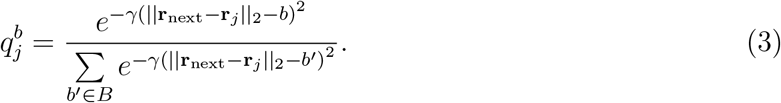

Here **r**_next_ is the ground truth position of the next atom, **r**_*j*_ is the position of an already placed atom, *N* is the number of preceding atoms, *γ* determines the width of the expansions, *B* are the 300 binned distances between 0 *Å* and 15 *Å*, and 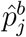 is the probability that the model assigns for the distance between **r**_*j*_ and **r**_next_ to fall into distance bin *b* ∈ *B* at the current step. In steps where the ground truth type is the stop type, the loss on distance predictions is set to zero as no distances are predicted. Descriptions about the hyper-parameters used in this work is provided in SI.

### Training data

Therapeutic candidates interact with target proteins either by forming a covalent bond or non-covalently with non bonding interactions. Depending on the kind of interaction, they are known as covalent or non-covalent drug candidates. The focus of our study is to develop a general framework capable of producing both covalent and non-covalent novel therapeutic candidates with specific scaffolds, so we performed experiments for two different datasets.

First, we performed experiments on covalent inhibitors data (hereafter called covalent dataset) taken from multiple sources.^32,33^ For the covalent dataset, we used ∼4000 candidates from a database of FDA approved drugs^32^ and cysteine molecules from the enamine database^33^ with 6 different scaffolds namely acrylamides, chloroamides, nitriles, disulfides, maleimides, and pyrodines.^32^ These functional groups react with the cysteine residue of the target protein by forming covalent bonds. The distribution of each scaffold in the data set is provided in the pie chart in Figure 2. Nearly 95 % of the training set is dominated by 3 scaffolds. We later show that, irrespective of the fraction of data for each scaffold, our model generalize equally well for all of them. SMILES strings of the molecules are extracted from the respective databases. RDkit^34^ with MMFF94^35^ forcefield was used to convert SMILES into the 3D coordinates required as an input for our model.

**Figure 2:**
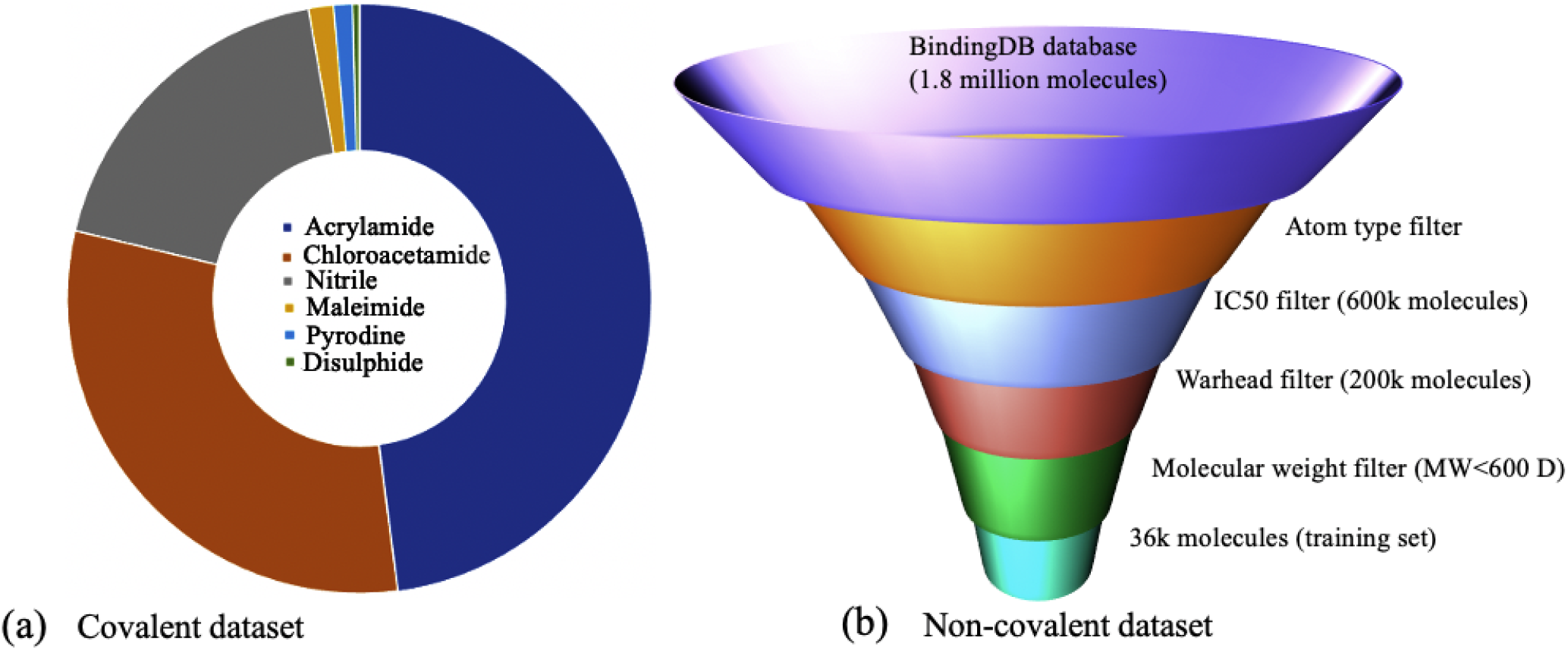
(a) Distribution of covalent dataset based on scaffolds. (b) Filtering criteron used to generate non-covalent training dataset. See SI for details used for atomtype and warhead filter.

In addition, for non-covalent inhibitor design we curated and filtered a large dataset of synthesizable molecules from BindingDB, ^36^ MCULE^37^ and Enamine^33^ databases to create the non-covalent dataset. We used different filtering criterons as shown in Figure 2 for creating the dataset. Our non-covalent inhibitor design model is trained with 36k molecules consisting of 10k unique scaffolds. For the non-covalent dataset, we use Murcko scaffolds^24^ as a definition of scaffolds, which demonstrates the flexibility of our model not only in allowing different scaffold definitions, but also for generating non-covalent inhibitors. We used RDkit to obtain Murcko scaffolds from SMILES strings of molecules in the training set. For generation with this dataset, we randomly select 25 out of the 10k scaffolds and generate 1000 molecules for each of them, providing ample generated molecules to assess the performance of the model.

## Results and Discussion

Despite tremendous effort, COVID-19 lacks effective therapeutics. As of now, no antiviral drugs were developed against the closely related coronavirus, SARS-CoV-1 or MERS-CoV, regardless of previous zoonotic outbreaks.^38^ To identify starting points for such therapeutics, our focus is to develop a domain informed ML framework to generate covalent electrophiles and non-covalent inhibitor candidates against the SARS-CoV-2 main protease (Mpro) and SARS-CoV-2 non-structural protein endoribonuclease (NSP15), two main viral proteases essential for viral replication. Most of the therapeutic candidates for SARS-CoV-2 have been taken from existing databases to screen against the target proteins. However, it is challenging to generate novel yet target specific molecules knowing the functionality and scaffold that can lead to high potency and efficacy.

### Covalent antiviral inhibitor design for Mpro

Using the covalent antiviral dataset, we first trained the model to generate molecules with 6 different scaffolds that are common electrophilic warheads for different drug applications. For each of the scaffolds, we generated 2000 molecules and inspected them for their validity, uniqueness and novelty. To calculate the percentage of valid, unique, and novel molecules, we use

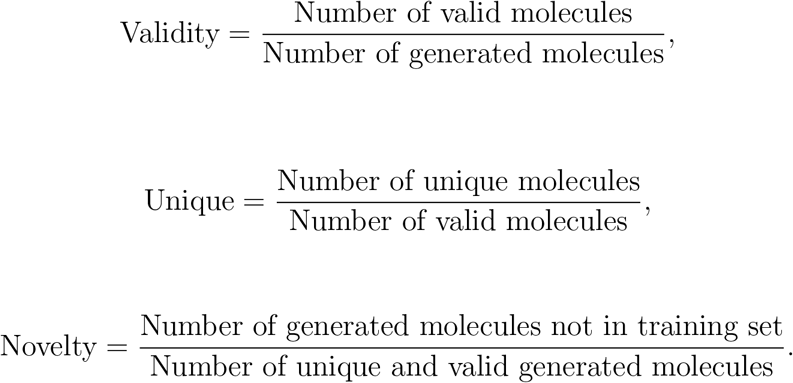

The validity of generated molecules is examined by converting generated 3D coordinates into canonical SMILES strings using the *xyz2mol* script from the Jensen group, ^39,40^ which relies on Rdkit.^34^ The conversion could also be done using only Rdkit or other open source tools like Open Babel but they are less reliable when determining bond orders during conversion. We then used the sanitize functionality of Rdkit to examine the validity of thus obtained SMILES strings. Alternatively, the validity of generated molecules can be measured by performing physics based simulations such as density functional theory. But due to enormous computational cost required to perform such calculations on thousands of generated molecules, we resort to empirical approaches for the same. To examine the novelty of generated molecules, we compared the Rdkit topological fingerprint similarity of the molecules in training set and generated set. The uniqueness metric is determined similarly by using molecular fingerprints. In addition, to further authenticates the performance of our model in generating valid and synthesizable molecules, we also query the MCULE database^37^ for generated molecules to check how many already exist in the MCULE dataset. The performance of our model in terms of these metrics is listed in Table 2.

**Table 2:** Table showing the statistics of valid, unique, and novel molecules generated for different scaffolds. The number of generated molecules that exist in MCULE database (not in the training set) is also listed in column ‘Known’. For the model trained on the non-covalent dataset, mean values of validity, uniqueness and novelty for 25 different scaffolds is provided. For comparison, performances of recent methods from the literature are also provided. However, note that literature results stem from experiments with different datasets than the ones used in this work.

|  | Validity (%) | Uniqueness (%) | Novelty (%) | Known |
| --- | --- | --- | --- | --- |
| Scaffolds | Covalent dataset |  |  |  |
| Acrylamides | 79 | 96 | 99 | 59 |
| Chloroamides | 83 | 93 | 99 | 34 |
| Pyrodines | 84 | 83 | 100 | 71 |
| Maleamides | 86 | 85 | 99 | 73 |
| Nitriles | 81 | 97 | 100 | 59 |
| Disulphides | 75 | 98 | 100 | 1 |
| Piperazine<sup>a</sup> | 80 | 92 | 100 | 52 |
|  | Non-covalent dataset |  |  |  |
|  | 90 | 73 | 100 |  |
|  | Literature |  |  |  |
| G-SchNet<sup>26</sup> | 77 | 92 | 88 | — |
| Lim et al.<sup>21</sup> | 99 | 85 | 99 | — |
| DeepScaffold<sup>23</sup> | 99 | 69 | — | — |
| GraphVAE<sup>41</sup> | 56 | 76 | 62 | — |
| MolGAN<sup>41</sup> | 98 | 10 | 94 | — |

<sup>a</sup> Novel scaffold

The performance of our model is similar to existing scaffold-based generative models in terms of generating valid, unique and novel molecules. For all the scaffolds in the covalent dataset, our model performs similarly well, with on average 92% uniqueness among the generated molecules. 81% of generated molecules are valid and ∼100% are novel. These metrics remain similar even for the molecules generated using a novel scaffold (piperazine) as starting point, thus demonstrating the transferrability of our model to scaffolds not in the training set. Compared to the existing generative models in the literature, our model shows superior performance in generating unique and novel molecules, while the percentage of valid molecules generated is in general slightly lower than for other generative models. We however note that these models were trained on different datasets, making a direct comparison of the reported numbers difficult. Moreover, the performance of our model is especially promising when one takes into account the relatively small amount of training data used (4000) compared to cited models from the literature which were trained on larger training sets. Training our model on larger training sets might further improve the reported statistics. In addition, compared to ours, models from the literature were trained to generate relatively small molecules with the QM9 dataset. When querying the MCULE database, we found that some of the molecules generated for each scaffold are already known and available in the database, demonstrating the success of our model in generating synthesizable molecules. This also holds for the molecules generated with the novel scaffold piperazines.

An important goal of our work is to generate novel molecules with drug-like properties while retaining desired scaffolds. To this end, we do not directly condition molecule generation on the desired properties but instead constrain it to the generation of molecules with desired scaffolds. We expect that this will indirectly constrain the properties, as well. The properties of interest are synthetic accessibility (SA) score, quantitative estimation of drug-likeliness (QED), and the partition coefficient (logP). The SA score measures the synthesizability of generated molecules and has values in the range 0-10, where the lower end suggests increased accessibility. QED is a useful measure for quantifying and ranking the drug-likeness of a compound. The values range from 0 for unfavorable to 1 for favorable molecules. The partition coefficient, logP, estimates the lipophilicity or hydrophilicity of a compound. It measures the physical nature of a compound and its permeability and ability to reach the target in the body. A positive logP value indicates the compound is lipophilic and a negative logP value indicates a hydrophilic compound.

We compare the properties of the generated molecules with the ones in the training set to see whether our model can generate new molecules with properties similar to those of the molecules in the training set. Ideally, having similar statistics of properties is an indicator that our model is performing as expected with the constraints imposed upon it. For the statistical analysis, we report the mean and standard deviation of the SA, LogP, and QED scores in both the training and the generated sets in Table 3. The mean SA score of both generated and training set molecules falls in the lower half of the SA scale 0−10, implying in general synthesizability of generated molecules. Slight deviation observed between the two sets can be attributed to the lack of explicit conditioning on target properties. The mean value of QED for generated molecules is slightly lower (on average by 0.2 units) compared to molecules from the training set. However, the model also generated molecules with high QED, i.e. strong drug-likeliness. logP follows similar trends for its mean value among two sets. We consistently observed relatively large standard deviation for SA, QED and logP in generated molecules for each scaffolds, reflecting diversity in generated molecules compared to the well curated training dataset. To further visualize this data, we display the probability density plots for SA, QED, and logP of the molecules in the training set and the generated set for each scaffolds in Figure 3. Solid lines mark the distributions of generated molecules while dashed lines correspond to molecules in the training set. The distributions of generated molecules with respect to the SA score in Figure 3(a) show that a good fraction of generated molecules are experimentally synthesizable. Moreover, the distribution of the SA score for the novel functional group, piperazine (not in training set), is similar to other scaffolds in the training set, showing the transferability of our model. This also demonstrate the success of our model in generating experimentally synthesizable molecules, which is a big issue with most generative models. For the logP metric (see Figure 3(c)), similar distributions are observed between generated and training molecules. We were able to generate both lipophilic as well as hydrophilic compounds as indicated by positive and negative logP, respectively, with the former category being the majority, similar to the molecules in the training set. This again indicates that our model is generating novel molecules with properties similar to the training set. From the QED distribution plot (Figure 3(b)), we see that the majority 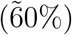 of the molecules have a QED score greater than 0.5, with a good chunk of molecules being close to 1 as evident from the peaks of probability distribution curves around 0.9. Minor discrepancies between the properties of generated molecules and the training set may be due to the lack of directly constrained property optimization in our work. Although our model generates molecules with desired properties, it would be interesting to see its performance when explicitly constraining the desired property range. However, this is beyond the scope of our current work and is kept aside for future work.

**Table 3:** Statistics of molecules from the training and generated data set, respectively for each scaffold. The mean and standard deviation for each property in each set are provided.

|  |  | Training Set |  |  | Generated Set |  |  |
| --- | --- | --- | --- | --- | --- | --- | --- |
|  |  | SA | LogP | QED | SA | LogP | QED |
| Chloroamides | Mean | 2.55 | 2.30 | 0.84 | 4.59 | 1.73 | 0.65 |
|  | Std | 0.57 | 1.00 | 0.15 | 1.31 | 1.97 | 0.23 |
| Acrylamides | Mean | 2.65 | 2.40 | 0.76 | 4.26 | 2.00 | 0.60 |
|  | Std | 0.55 | 1.33 | 0.21 | 1.42 | 1.99 | 0.25 |
| Disulphides | Mean | 2.94 | 2.64 | 0.88 | 5.60 | 3.15 | 0.52 |
|  | Std | 0.89 | 0.82 | 0.22 | 0.95 | 2.15 | 0.26 |
| Pyrodines | Mean | 2.90 | 1.62 | 0.70 | 4.62 | 0.80 | 0.52 |
|  | Std | 0.59 | 1.47 | 0.25 | 1.55 | 1.71 | 0.28 |
| Maleamides | Mean | 2.30 | 1.32 | 0.66 | 4.36 | 0.72 | 0.63 |
|  | Std | 0.27 | 1.02 | 0.17 | 1.40 | 1.62 | 0.24 |
| Nitriles | Mean | 2.40 | 3.12 | 0.87 | 4.47 | 2.10 | 0.61 |
|  | Std | 0.40 | 1.00 | 0.13 | 1.28 | 1.84 | 0.23 |
| Piperazine<sup>a</sup> | Mean | — | — | — | 4.75 | 1.09 | 0.54 |
|  | Std | — | — | — | 1.31 | 1.86 | 0.29 |

<sup>a</sup> Novel scaffold

**Figure 3:**
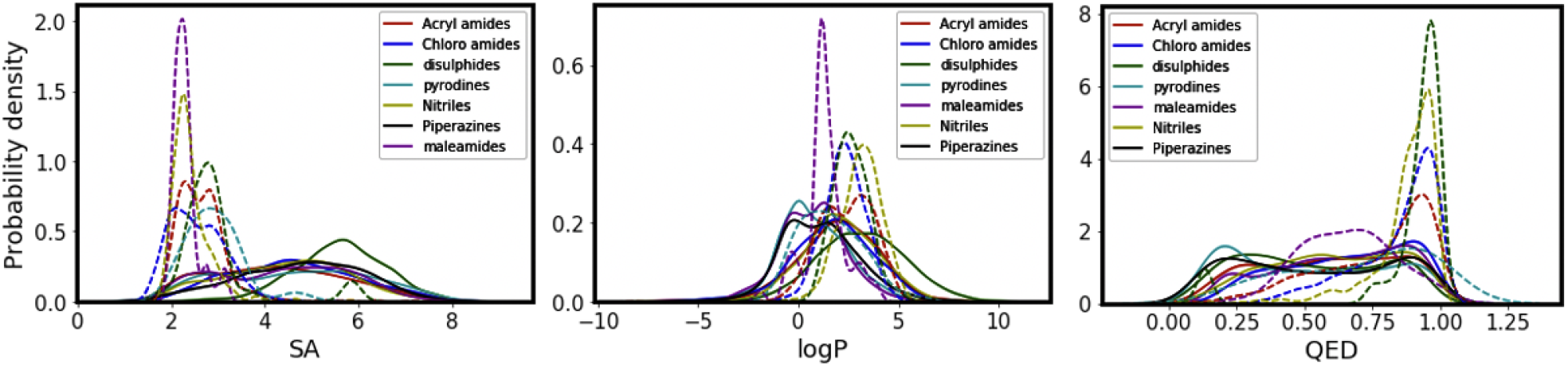
Probability density plots of SA score, logP, and QED for molecules in the training set as well as generated set for each functional group. Solid lines correspond to metrics of data in the generated set, whereas dashed lines of same color correspond to molecules in the training set.

We further analyzed the diversity of molecules using heatmaps of the Tanimoto coefficient between molecules within the training set (Figure 4(a)) and within the generated set (Figure 4(b)). The tanimoto coefficient is a measure of the similarity of molecules. The heatmap shows that the training set we use is quite diverse as evident by the many green spots (low similarity). A similar heatmap is observed for the generated set, showing that generated molecules are quite different from each other, while predominantly maintaining similar properties (as discussed before). We also note that our model generates diverse molecules in terms of their size, i.e. the number of atoms, while always preserving the given scaffolds.

**Figure 4:**
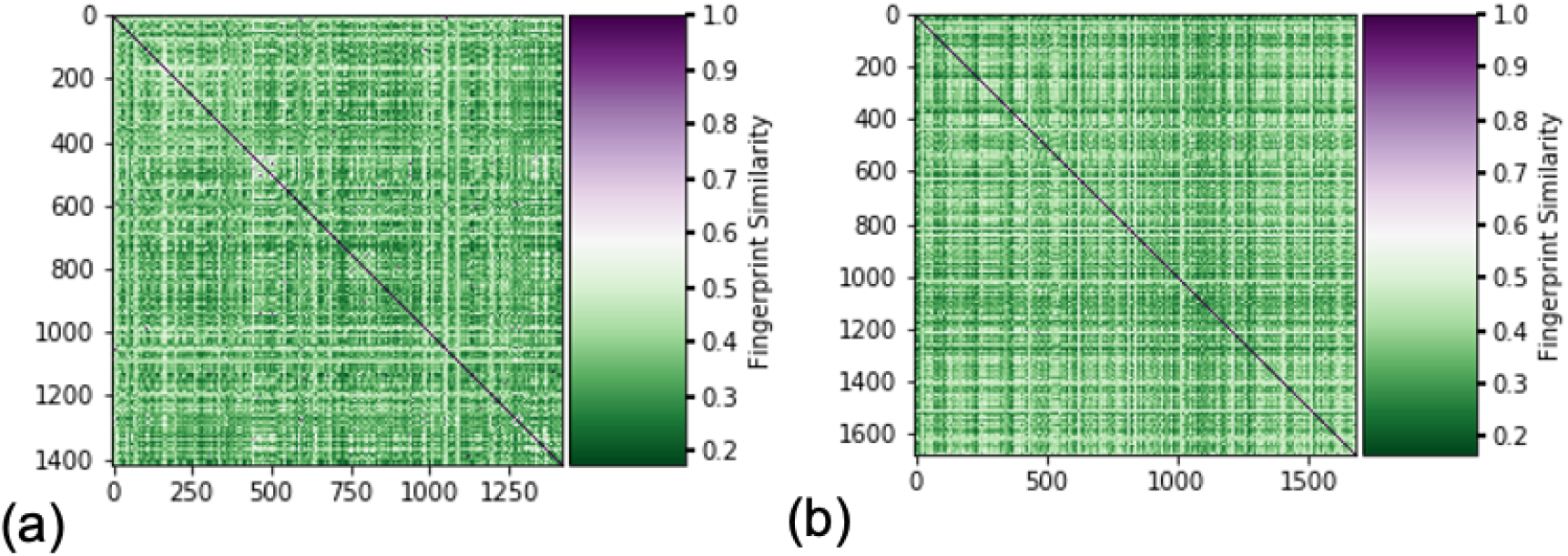
Heatmap showing the fingerpoint similarity between molecules in the training set and the generated set (b) for the scaffold acrylamide.

To check the transferability of our model to generate valid molecules for functional groups that are not in training set, we generated 2000 molecules with “Piperazine” as the starting building block. Generated molecules are checked against the MCULE databases to see if any of the generated molecules are already known. We found that nearly 50 of the molecules generated are available in the MCULE database. This shows the capacity of our model to generate valid, synthesizable molecules even for novel scaffolds. The distribution plot for the SA, logP, and QED of the molecules generated for piperazine is included in Figure 3. It shows that the properties of molecules generated follow similar distributions as for other functional groups.

We visualize representative generated molecules that we also found in the MCULE database with corresponding SA score, QED, and logP values along with the corresponding MCULE ids in Figure 5. Overall, our results show that our model constrained to generating molecules with desired scaffolds indirectly also successfully constrains the properties. Despite the significant variation in the amount of training data for each scaffold, our model consistently generates valid, unique, novel, and experimentally synthesizable molecules with desired drug-like properties for each scaffold within and outside of the training set.

**Figure 5:**
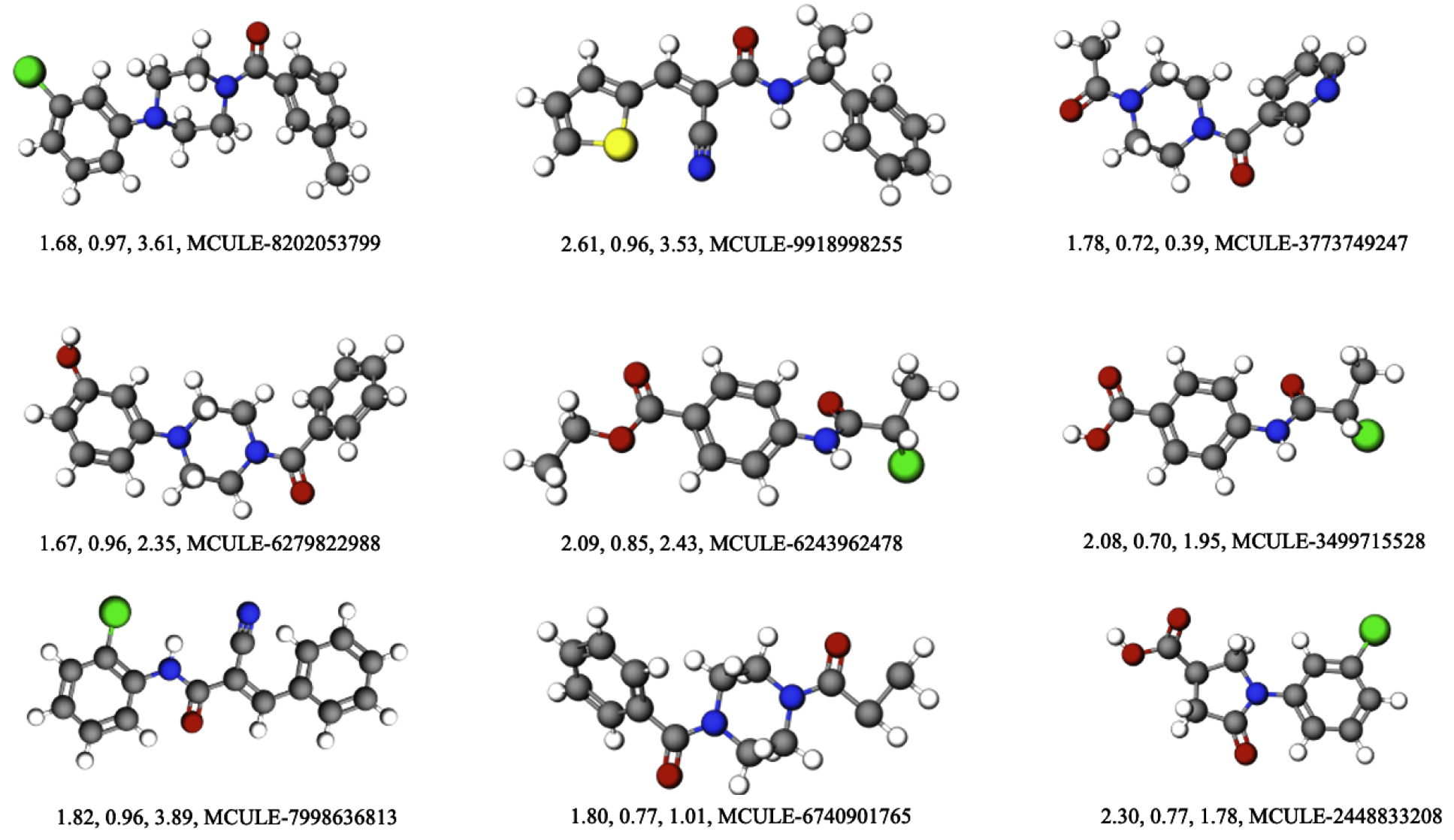
Sample of generated candidates along with their SA score, QED, logP values, and corresponding MCULE ids. These candidates are synthesizable and available to order from MCULE database.

### Binding affinities of covalent inhibitors against Mpro

Finally, as a proof of concept application for generated molecules, we docked them against main protease (Mpro). Mpro is the key enzyme of SARS-CoV-2 that gets the maximum attention because of its ability to trigger viral replication and transcription. Significant effort has been made since the rapid rise of SARS-CoV-2 worldwide to find therapeutic candidates/vaccines that have desired activity against its protein. Most of the early efforts were focused on drug repurposing using already known drug molecules. For future pandemic events, it is possible that an effective drug molecule for repurposing is not yet known. In those scenarios, models that generate novel molecules with certain functionalities as proposed here will be an efficient alternative. First step for such generated molecules is to examine their efficacy against the target protein using docking simulations. Docking simulations use empirical approaches to determine the favorable/unfavorable binding of ligands with target proteins and numerically rank them using a docking score.

For our molecular docking simulations we utilized the AutoDock for Flexible Receptors (ADFR) package.^42^ Ligands were covalently bound to Cystine-145 of the target protein (PDB ID: 6WQF), which is part of a catalytic dyad formed with Hystine-41. We compared the docking score of generated molecules against the training molecules in the covalent dataset which is shown in Figure 6. A larger magnitude of the docking score implies higher favorability for the docking process. We found that generated molecules show similar docking performance when compared to the molecules in the training covalent dataset as illustrated in the violin plots and the corresponding mean docking score noted in the labels of the x-axis. For the majority of scaffolds, including the novel scaffold piperazines and the three scaffolds that make up 95% of the training data, the generated molecules on average show higher affinity for docking against the Mpro-target protein than molecules in the covalent dataset. The only scaffolds that have a smaller mean docking score compared to the training molecules are maleamides and pyrodines with docking scores of 8.96 and 8.64, respectively.

**Figure 6:**
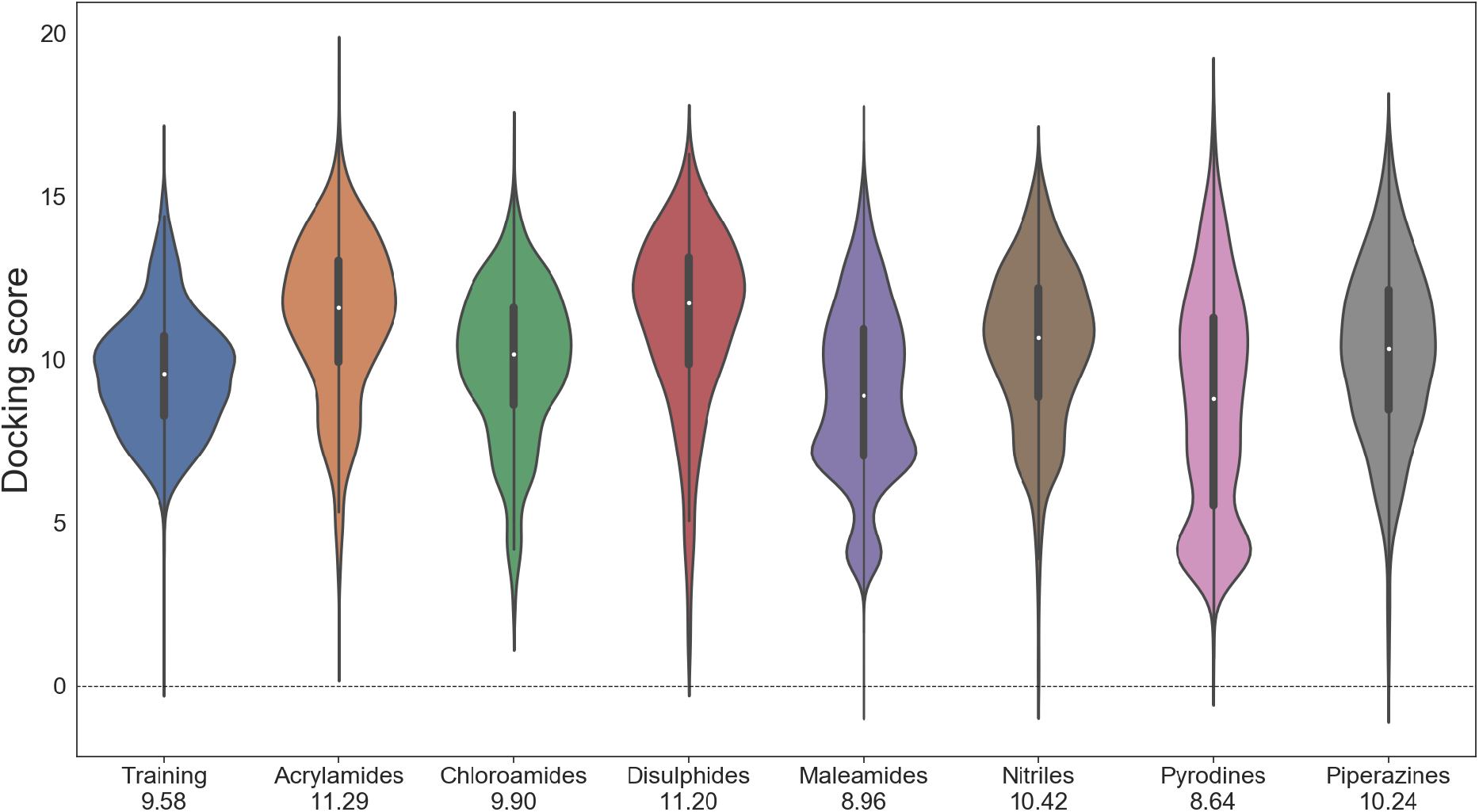
Violin plots showing the distribution of the docking score against the MPro protein for generated molecules with different scaffolds and training molecules in the covalent dataset. Larger values imply favorable binding.

### Non-covalent antiviral inhibitor design for NSP15

With the goal of generating non-covalent inhibitors for SAR-Cov-2 targets, we trained our model on the non-covalent dataset using Murcko scaffolds. The training data consist of 36k molecules with 10k unique scaffolds. The performance of our model trained for generating non-covalent inhibitors is similar (Table 2) to the one for the covalent dataset in terms of validity and novelty. However, the percentage of unique molecules generated drops to a mean value of 73 % for about 25 different scaffolds. This may be a direct consequence of the limited number of molecules (on average 4) for each scaffold in the non-covalent training set. When generating 1000 molecules for each of the 25 scaffolds, the model repeats some of the generated molecules. However, the absolute amount of uniquely generated molecules per scaffold is still remarkable considering the limited number of training examples per scaffold.

As a part of DOE National Virtual Biotechnology Laboratories (NVBL) therapeutic design project, we screened millions of compounds in repurposing libraries of drug compound for activity against nsp1-nsp15 from SARS-CoV-2 followed by experimental validation. In particular, the coronavirus nonstructural protein NSP15 is highly conserved among coronaviruses. It is also a key component for viral replication with no corresponding counterpart in host cells which makes it an intriguing candidate for drug development. Our recent computational and experimental results demonstrated that Exebryl-1, a ß-amyloid anti-aggregation molecule designed for Alzheimer’s disease therapy can bind to NSP15 but it did not have sufficient anti-viral activity in cell-based assays for immediate drug repurposing efforts. ^43^ This provide us an interesting target to optimize the Exebryl-1 hit based on 3D-Scaffold framework with better activity and antiviral properties. Our goal is to lead optimization together with in silico molecular docking calculations onto the crystal structure of NSP15.

As a test case example, we generated non-covalent inhibitors for the SAR-CoV-2 non-structural protein endoribonuclease (NSP15) target (PDB ID: 6XDH) by optimizing Exebryl-1 based compounds.^43^ Exebryl-1 has experimentally been found to be active ^43^ against NSP15 from high-throughput assay screening from drug and lead repurposing libraries. Our goal is to modify and generate more active compounds against the NSP15 target by building molecules on top of Murcko scaffolds of the Exebryl-1 molecule. When examining the structure-activity relationship, some of such generated molecules (see Figure 7) show good binding-activity against the NSP15 target. Moreover, these molecules are easily synthesizable (low SA scores) and have desired drug likeliness (large QED values). Generated molecules from our work that showed high activity against NSP15 from docking and molecular dynamics simulations are further being investigated by our experimental collaborators.

**Figure 7:**
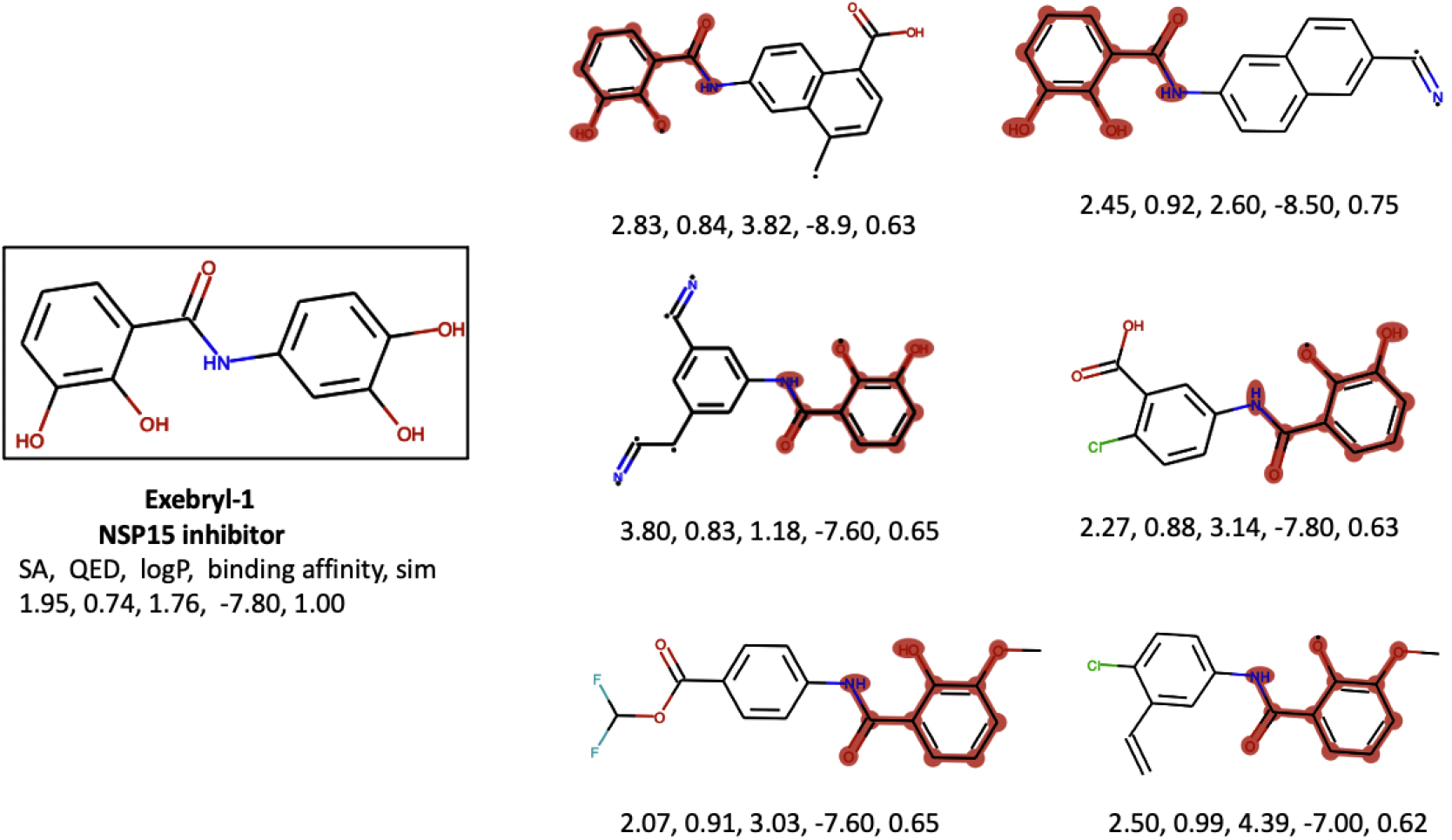
Exebryl-1 and representative generated molecules from our 3D-Scaffold framework with high binding affinity against NSP15 protein-target. For each molecule, we list the SA score, QED, logP, binding affinity, and fingerprint similarity (labelled sim in figure) with respect to experimentally known NSP15 inhibitor Exebryl-1. The scaffold used for optimization is highlighted in red in generated molecules.

## Conclusions

In this report, we developed a generative framework that can generate 3D coordinates of therapeutic candidates with any desired scaffold. The model is trained end-to-end incorporating robust atomistic representation learning techniques and generates 3D coordinates from the learned probability distributions of atom types and the pairwise distances. Due to starting the sequential atom-by-atom generation scheme of our framework from a given scaffold, the desired scaffold is 100 % guaranteed in the generated 3D coordinates. We use covalent and non-covalent antiviral datasets to optimally narrow the search towards novel compounds with therapeutic significance that are reasonable to design as covalent and non-covalent inhibitors. We show that our model generates predominantly valid, unique, and novel molecules that have therapeutic drug-like properties similar to the molecules in the training set. The success of our framework lies in generating synthesizable molecules with desired properties without directly constraining on the target properties. Moreover, it performs well for relatively small volumes of training data and generalizes equally well for generating molecules with a new scaffold, which demonstrates the transferability of the proposed framework.

Our framework offers the advantage that the generated 3D-coordinates of molecules can be directly used for further simulations such as DFT, MD, or docking calculations in contrast to SMILES or graph based models where empirical approaches are used to generate 3D coordinates. As an application, the 3D coordinates of generated molecules from our work were examined for their interaction against the Mpro and NSP15 targets of SARS-CoV-2 using docking simulations. Our results show that generated molecules have favorable interaction against the target protein similar to the molecules in the training set. This holds true for novel scaffolds as well. Although we used our framework to generate covalent and non-covalent inhibitors in this work, our model in principle can be used to generates any kind of molecules with desired scaffolds making it applicable to many domains. We believe that the robust performance of our model on relatively small data sets and its generalization on new scaffolds provides an efficient and flexible way of generating new molecules while simultaneously optimizing the functionalities by constraining the types of scaffolds included. Further improvement in the performance of the 3D-Scaffold framework may be observed by generating molecules while also explicitly constraining on the target properties or by generating molecules with more than one critical scaffold.

## Acknowledgement

This research was supported by the DOE Office of Science through the National Virtual Biotechnology Laboratory, a consortium of DOE national laboratories focused on response to COVID-19, with funding provided by the Coronavirus CARES Act. Pacific Northwest National Laboratory (PNNL) is a multiprogram national laboratory operated by Battelle for the DOE under Contract DE-AC05-76RLO 1830. Computing resources was supported by the Intramural program at the William R. Wiley Environmental Molecular Sciences Laboratory (EMSL; grid.436923.9), a DOE Office of Science User Facility sponsored by the Office of Biological and Environmental Research and operated under Contract No. DE-AC05-76RL01830. We thank Darin Hauner at PNNL for discussion on the covalent docking simulations. Provisional application of small molecule candidates designed from this deep learning work is pending.

## Supporting Information Available

Codes used for this work are available at https://gitlab.pnnl.gov/computational_data_science/ml/3d_scaffolds.

## Notes

### Competing Interest Statement

The authors have declared no competing interest.

